# FootprintCharter: unsupervised detection and quantification of footprints in single molecule footprinting data

**DOI:** 10.1101/2025.03.31.646464

**Authors:** Guido Barzaghi, Arnaud R Krebs, Judith B Zaugg

## Abstract

**Summary:** Single molecule footprinting profiles the heterogeneity of TF occupancy at *cis*-regulatory elements across cell populations at unprecedented resolution. The single molecule nature of the data in principle allows for observing the footprint of individual transcription factors and nucleosomes. However, we currently lack algorithms to quantify these occupancy patterns of chromatin binding factors in an automated way and without prior assumptions on their genomic location. Here we present *FootprintCharter*, an unsupervised tool to detect and quantify footprints for transcription factors (TFs) and nucleosomes from single molecule footprinting data. *FootprintCharter* allows for the quantification of complex molecular states such as positioning of unphased nucleosomes and combinatorial co-binding of multiple TFs.

**Availability and implementation:** *FootprintCharter* is freely available on Bioconductor with version 2.1.6 of https://bioconductor.org/packages/SingleMoleculeFootprinting with the functions *FootprintCharter, PlotFootprints* and *Plot_FootprintCharter_SM*.

## 1 Introduction

Single molecule footprinting (SMF) maps DNA-protein interactions at *cis*-regulatory elements at single molecule resolution. It does so by marking cytosines that are not protected by DNA-bound proteins with exogenous methyl-transferases and by measuring those methylation marks with bisulfite sequencing (Krebs et al. 2017; Kleinendorst et al. 2021). Recent studies demonstrated the ability of SMF to measure the heterogeneity of transcription factor (TF) binding(Sönmezer et al. 2021) (Figure 1A), polymerase occupancy(Krebs et al. 2017; Chatsirisupachai et al. 2024) and chromatin accessibility(Kreibich et al. 2023; Baderna et al. 2025) at the single molecule level across cell populations. Yet, we are still lacking a tool for the automatic quantification of occupancy patterns of chromatin binding factors without prior assumptions on their genomic location such as mapped TF motifs or transcription start sites.

**Figure 1.**
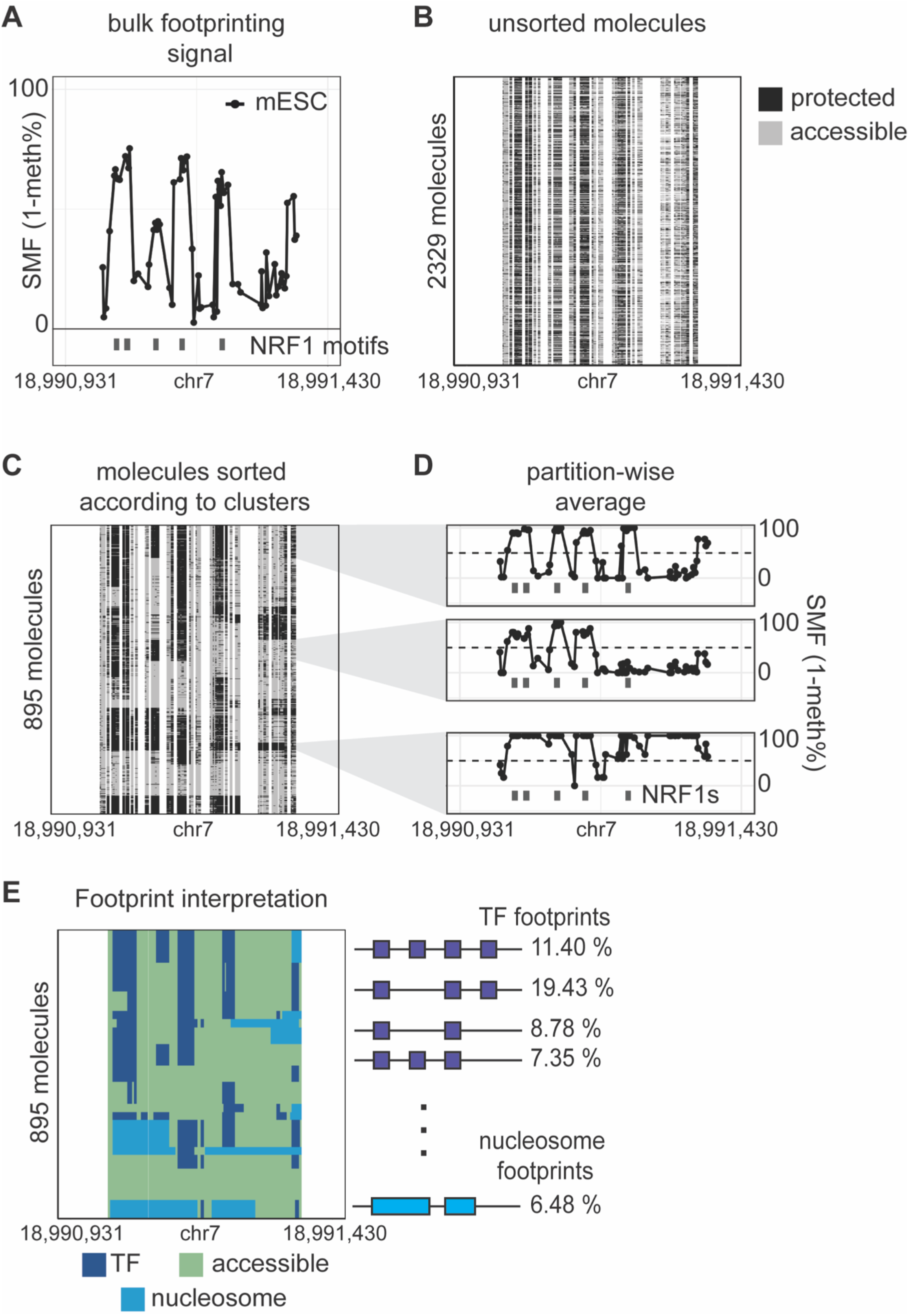
*FootprintCharter* allows for the unsupervised quantification of single molecule footprinting patterns at complex loci. (A) Example locus bound by NRF1 at five distinct motifs. The y-axis shows the average SMF signal (1-methylation %) at individual GpCs and CpGs, which can be interpreted as a measure of genomic occupancy frequency. Already the average SMF signal reveals short, but highly frequent, footprints overlapping the TF motifs. (B) Heatmap displaying the unsorted molecules covering the locus. Each column shows an individual genomic cytosine and each row a single molecule. Each single cytosine on a single molecule, is colored according to its binary occupancy status (accessible in grey, protected in black).(C) Single molecules sorted according to the unsupervised clustering results. (D) Cluster-wise average SMF signal used for footprint detection. Short footprints (5-75bp) are interpreted as TFs, footprints wider than 120bp are interpreted as nucleosomes. (E) Single molecules annotated with biological state by footprint detection results (TF in purple, accessible in green, nucleosome in blue).

Complementing the repertoire of computational tools for SMF(Kleinendorst et al. 2021; Doughty et al. 2024) and of unsupervised tools for single molecule genomics(Tullius et al. 2024; Vollger et al. 2024), we developed *FootprintCharter*, a novel unsupervised molecular classifier and footprint detection algorithm tailored for SMF. Differently from previous methods, *FootprintCharter* explicitly detects footprints for transcription factors (TFs) and nucleosomes and quantifies their frequency across cell populations. Notably, this tool performs these quantifications independently of orthogonal TF motif annotations. *FootprintCharter* allows for the estimation of frequencies for complex molecular patterns along single molecules, including various combinatorics of large TF clusters and nucleosome arrays. The *FootprintCharter* implementation is freely available as part of the *devel* version of our previously reported *SingleMoleculeFootprinting* R/Bioconductor package(Kleinendorst et al. 2021; Gentleman et al. 2004; Huber et al. 2015).

## 2 Implementation

### 2.1 Input

*FootprintCharter* takes as input single molecule methylation matrixes as produced by the function *CallContextMethylation* of our *SingleMoleculeFootprinting* R/Bioconductor package (Figure 1B). Internally, methylation calling at the single molecule level is carried using the Bioconductor package *QuasR*(Gaidatzis et al. 2015). The *QuasR* output is converted to sparse matrixes using the *Matrix* R package(Bates, Maechler, and Jagan 2000).

*FootprintCharter* quantifies footprints independently of prior TF motif annotations. Therefore, it focuses on user-defined genomic intervals encoded as *GRanges* objects from the *GenomicRanges* R/Bioconductor package (Lawrence et al. 2013). Read length should be considered when defining these windows. For instance, we experienced that 80bps is a suitable interval width for 150 paired-end SMF data.

### 2.2 Unsupervised clustering of molecules

First, binary methylation values are smoothed along single molecules by computing, at each genomic position, a rolling mean over a 40bp window. Secondly, a matrix of pairwise Euclidean distances among single molecules is computed using the cytosine methylation signal along molecules as features. This matrix is computed with the *parDist* function of the *parallelDist* R package. Unsupervised clustering is performed on the resulting matrix using the partitioning around medoids algorithm (pam), implemented in the *pam* function of the *cluster* R package(Schubert and Rousseeuw 2021) (Figure 1C). The number of clusters (*k*) is initially set by the user and iteratively reduced by *FootprintCharter* until all clusters are populated by at least *n* molecules, where *n* is also user-defined. This ensures robust detection of footprints in the next step. Default values are set at *k*=16 and *n*=5, which have empirically worked well in our hands for SMF datasets produced with a bait-capture enrichment step(Kleinendorst et al. 2021; Sönmezer et al. 2021).

### 2.3 Footprint detection

For each cluster, which corresponds to a biologically distinct chromatin occupancy pattern, *FootprintCharter* computes the average SMF signal at each cytosine using the input binary methylation values. Within those average tracks, cytosines are defined as part of footprints if their resulting median SMF signal exceeds 50% (Figure 1D). Consecutive stretches of footprinted cytosines that are between 5 and 75 bp in width are interpreted as TF footprints. Stretches longer than 120bp are interpreted as nucleosome footprints. Footprints are considered as “unrecognized” if they are not flanked by accessible cytosines on both sides since their full width cannot be established, as it happens at the edge of molecules (Figure 1E).

### 2.4 TF footprint aggregation and annotation

*FootprintCharter* detects footprints separately for each cluster. However, measuring TF-DNA interactions is associated with a certain level of biological and technical noise. We for instance observed that the same protein-DNA contact might result in footprints that are partially shifted along different molecules. To account for this when estimating the frequency of footprints across cells, we aggregate TF footprints when they overlap by at least 75% of their width.

Users can also provide TF motif annotations, such as those from the JASPAR database(Mathelier et al. 2016), to label TF footprints. If provided, this annotation will also be used to aggregate TF footprints.

### 2.5 Output and data visualization

The steps above are performed by the *FootprintCharter* function. The output is a data frame reporting the coordinates and frequencies of the TF and nucleosome footprints detected around the input genomic interval.

The data transformations performed by *FootprintCharter* can be visualized using the functions *PlotFootprints* and *Plot_FootprintCharter_SM*.

### 2.6 Note on generalization and future developments

Albeit specific to SMF, *FootprintCharter* does not make assumptions on the footprinted context and it could work with data that include adenosine methylation such as Fiber-seq(Stergachis et al. 2020) or SMAC-seq(Shipony et al. 2020). However, because the running time of the initial smoothing step grows linearly with the molecule length, further tool development might be necessary to exclude this step.

*FootprintCharter* is intended as an unsupervised algorithm, but it still relies on orthogonal motif annotations to label TF footprints. A further development could include the *de novo* discovery of motifs from TF footprints aimed at labelling the orphan footprints discovered by the tool.

## Author contributions

Guido Barzaghi (Conceptualization [equal], Data curation [equal], Formal analysis [lead], Investigation [equal], Methodology [equal], Software [lead], Validation [lead], Visualization [lead], Writing – original draft [lead], Writing – review & editing [equal]). Arnaud R Krebs (Conceptualization [equal], Data curation [equal], Funding acquisition [lead], Investigation [equal], Methodology [equal], Project administration [equal], Resources [equal], Supervision [equal], Writing – original draft [supporting], Writing – review & editing [equal]). Judith B Zaugg (Conceptualization [equal], Funding acquisition [supporting], Methodology [equal], Project administration [equal], Resources [equal], Supervision [equal], Writing – original draft [supporting], Writing – review & editing [equal]).

## Acknowledgements

We acknowledge Christian Arnold, Maksim Kholmatov, Frosina Stojanovska, Aryan Kamal for helpful discussions during the development of *FootprintCharter*. We acknowledge Maksim Kholmatov and Jonas Oefelein for proofreading this manuscript.

## Data availability statement

The data used in this study was obtained from (Sönmezer et al. 2021) under ArrayExpress accession number E-MTAB-9033. The code used to generate the figure is available at https://github.com/Krebslabrep/Barzaghi-et-al_application-note.git.

